# Lesions Involving Medial Anterior Forebrain Pathway Circuitry Destabilize Phrase Timing in Adult Canary Song

**DOI:** 10.64898/2026.07.11.737998

**Authors:** M.R. Hulsey-Vincent, G. Vengrovski, E. Sova, T.J. Gardner

## Abstract

Basal ganglia-thalamocortical circuits are essential for learning complex motor sequences, yet how they control flexible motor behavior remains poorly understood. The homologous songbird Anterior Forebrain Pathway (AFP) drives song motor learning. Early lesion studies showed no effect on adult song, so the AFP was thought to be unnecessary for adult song performance. Recent evidence shows that the AFP influences song syntax in some species, revising this perspective. We revisit this question in adult canaries by performing bilateral excitotoxic lesions targeting the AFP’s lateral and medial subdivisions.

To measure behavioral changes, we developed a high-throughput song-annotation pipeline that adds a supervised classifier to the self-supervised TweetyBERT model. This removed the memory bottleneck of UMAP clustering, enabling phrase-level analysis across thousands of songs per bird. We find that lesions involving the medial AFP produce stuttering-like behavior, defined here as prolonged, variable syllable repetition before transitions, which significantly increases phrase-duration variability. This effect appeared only in birds with medial and lateral AFP lesions, not in birds with lateral-only lesions. This increased variability persisted throughout the post-lesion recording period and was accompanied by small changes in syllable acoustic structure.

Our results implicate the medial AFP in the ongoing control of phrase duration in adult canary song, challenging the view that the AFP is dispensable once song is learned. These findings position the medial AFP as a tractable model for understanding how basal ganglia and cortical dynamics maintain complex learned motor sequences.

**Significance Statement:** Basal ganglia-thalamocortical circuits are critical for learning complex motor sequences, but their role in performing learned behaviors remains unclear. We found that in adult male canaries, lesions involving the medial and lateral portions of the songbird Anterior Forebrain Pathway (AFP) resulted in an abnormal stuttering-like behavior. We did not detect comparable changes after lateral-only lesions, implicating circuitry involving the medial pathway, although we cannot isolate its independent contribution. These findings challenge the view that the AFP becomes unnecessary once canary song is learned or produced during the more stable breeding-season song, and offer a model for studying basal ganglia contributions to learned motor sequencing. We also develop a scalable annotation pipeline enabling analysis of thousands of songs per bird.

## Introduction

Complex motor skills, such as speaking, require combining simpler motor elements into flexible sequences. For example, saying the word “canary” requires planning a sequence of phonemes and coordinating vocal organs to produce each sound in sequenced order. How these sequences are planned and executed remains a central question in neuroscience (Lashley, 1951; Diedrichsen et al., 2026). Damage to the brain regions responsible for motor planning contributes to motor disorders such as Parkinson’s disease and speech disorders like stuttering (Craig-McQuaide et al., 2014).

Birdsong is a useful model for studying how these brain regions plan and execute action sequences. Songbird neural circuitry has simpler connections than mammalian cortical motor systems, but contains analogous motor control regions and cell types (Reiner et al., 2004; Jarvis, 2019). Canary song is built from syllables that the bird repeats in a “phrase” for about one second before switching to the next syllable in the song. These sequences have long-range order: what the bird sings at one point can influence what it may sing up to five to seven phrases later (Markowitz et al., 2013). Because songbird species vary in song complexity, songbirds are a useful model to study how brain regions orchestrate motor sequences at different levels of complexity and at different scales: syllable, phrase, and song (Fig. 2B).

**Figure 1.**
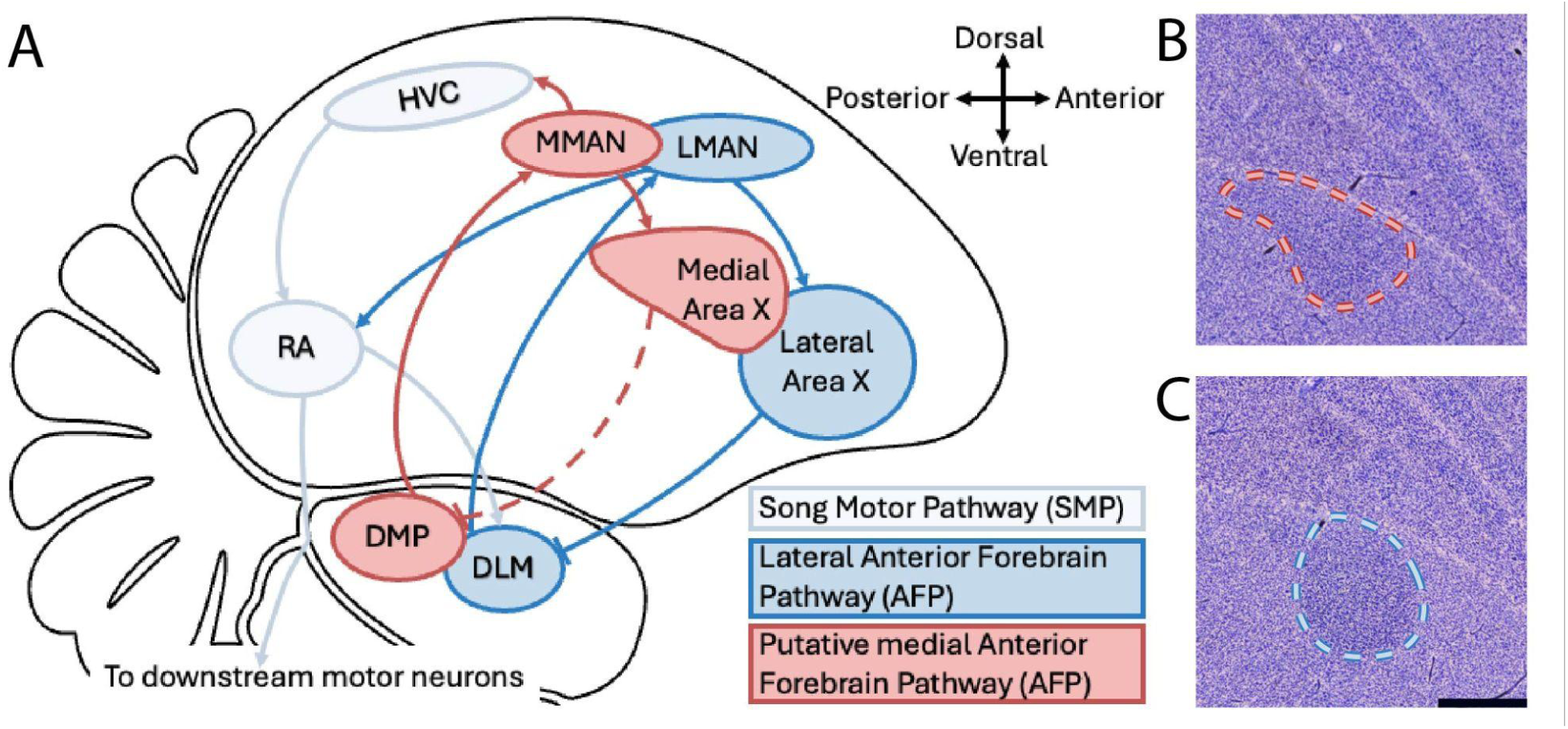
Anatomical reference for the medial and lateral Anterior Forebrain Pathway (AFP) targeting. A) Sagittal map of the organization of the AFP and the Song Motor Pathway. Avian brain outline adapted from (Nottebohm, 2005). The lateral AFP is shown in blue, and the medial AFP is shown in red. B) Sagittal Nissl-stained section showing intact medial Area X outlined by a red dashed line. C) Sagittal Nissl-stained section of lateral Area X, outlined by a blue dashed line. The scale bar indicates 500 μm. We divided Area X into medial and lateral portions based on the contours of the nucleus in the canary brain atlas (Vellema et al., 2011).

**Figure 2.**
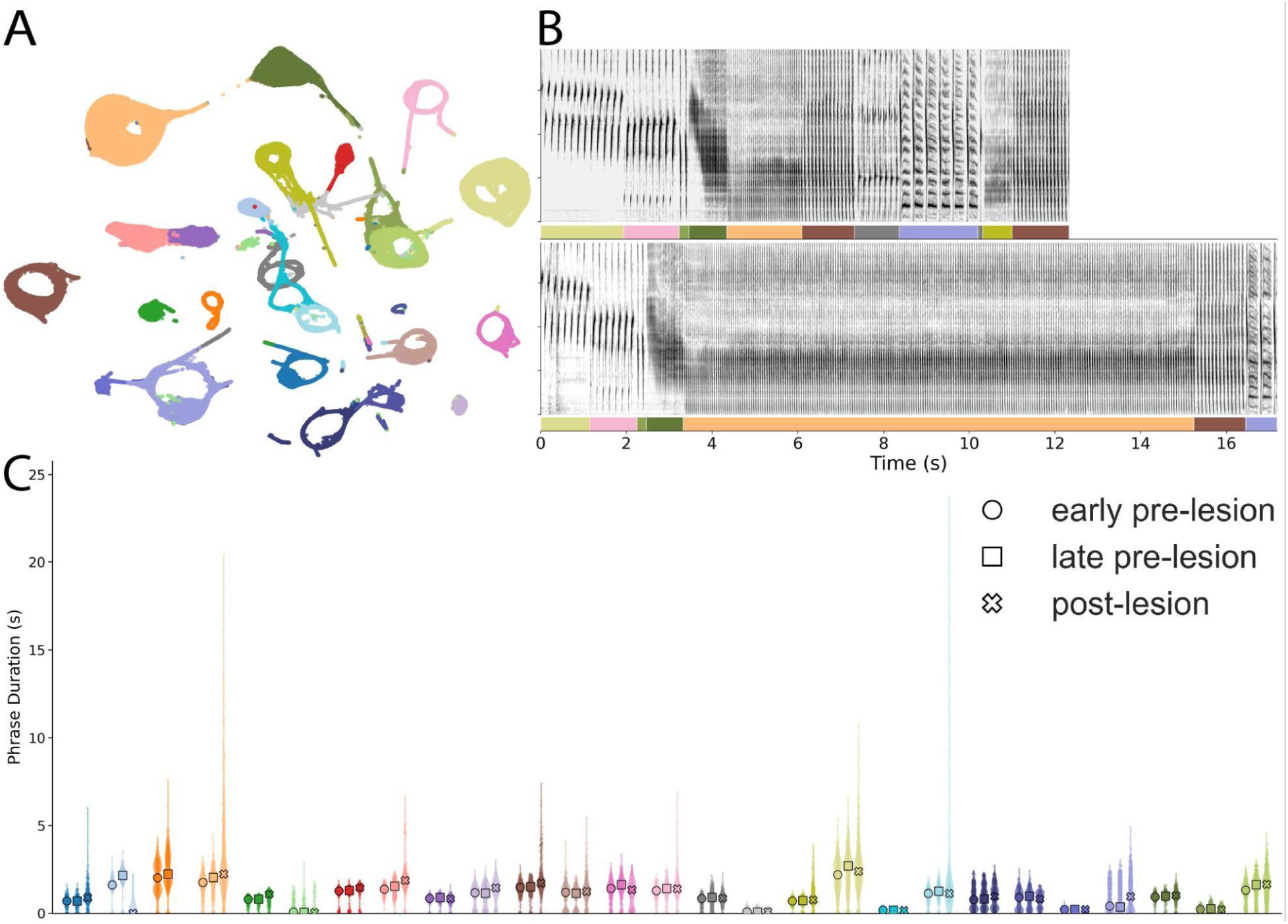
Lesion-associated stuttering-like syllable repetition in one canary. A) TweetyBERT embedding space from one canary (Bird 13), with clusters corresponding to syllable classes. Paired pre-lesion and post-lesion UMAPs for the bird are shown in Supplemental Fig. 2A. B) Representative pre-lesion song bout and post-lesion song from the same bird showing prolonged repetition of one syllable. Here, “stuttering-like” is used descriptively to refer to prolonged and variable syllable repetition before transition. Colored bars indicate TweetyBERT syllable annotations. C) Phrase duration summary across syllable labels in early pre-lesion, late pre-lesion, and post-lesion periods. A subset of syllables showed increased post-lesion phrase duration variability.

Birdsong is a fundamental model for studying the reinforcement learning of complex motor sequences. Song learning is driven by nuclei in the lateral portion of the Anterior Forebrain Pathway (AFP) (Fig. 1A): the cortical nucleus LMAN introduces variability that allows motor exploration, and the basal ganglia nucleus Area X biases that variability toward improved performance (Fee and Goldberg, 2011). Most research has focused primarily on the lateral AFP. However, the AFP also contains a parallel medial pathway comprising the cortical nucleus MMAN (magnocellular nucleus of the anterior nidopallium), the medial portion of Area X, and the thalamic nucleus DLM (Fig. 1A) (Nottebohm et al., 1976; Kubikova et al., 2007). This study examines the potential role of the medial AFP in adult song, particularly in timing and sequencing.

Early studies suggested the AFP was unnecessary in adult song, because lesions had no clear impact in zebra finches (Bottjer et al., 1984; Sohrabji et al., 1990) or canaries (Nottebohm et al., 1976). More recent work has challenged this view. In adult zebra finches that already repeated syllables at the end of their songs before the lesion, lesions of lateral Area X produced stuttering (Kubikova et al., 2014). In Bengalese finches, partial Area X lesions produced a transient stuttering-like behavior that eventually resolved and was not clearly related to lesion size (Kobayashi et al., 2001). Additionally, in Bengalese finches, MMAN lesions increased variability in syllable repeat number and increased uncertainty in the order of syllables (Koparkar et al., 2024). Together, these results suggest that the AFP remains involved in controlling adult song.

Our experiments build on these findings by performing bilateral lesions to the lateral and medial portions of the AFP in adult canaries during their breeding season. To quantify behavioral changes across the large number of songs produced, we developed a high-throughput annotation pipeline built on TweetyBERT, a self-supervised model that clusters canary syllables with high accuracy (Vengrovski et al., 2026). This pipeline overcomes the scalability limitations of Uniform Manifold Approximation and Projection (UMAP) and enables analysis of thousands of songs per bird. We found that lesions involving the medial AFP circuitry resulted in a stuttering-like behavior, characterized by an increase in phrase duration variability of select syllables, along with subtle changes in the acoustic structure of those same syllables. Neither effect appeared after lateral-only lesions or sham surgery. These findings suggest that the medial AFP continues to influence syllable transitions between repeated syllables in adult canary song, positioning the medial basal ganglia-thalamocortical pathway as a valuable model for understanding basal ganglia and cortical dynamics in motor sequencing.

## Materials and Methods Experimental Design

### Subjects, housing, and husbandry

Subjects were adult male canaries (medial+lateral AFP lesions n=9 [complete lesion n = 6, partial lesion n=3], lateral-only AFP lesions n=8 all partial lesions, sham controls n=4), age > 1 year; see Supplemental Table 1, obtained from Maryland Exotic Birds (Pasadena, MD). While in holding, birds were housed in flight cages on a light cycle synchronized to daylight hours in Eugene, Oregon, and given food (Kaytee Supreme Seed, Lafeber’s Canary Premium Daily Pellets), fresh water, and high-calcium grit (Kaytee Hi-Calcium Grit Supplement) *ad libitum*.

Housing and husbandry were performed by the University of Oregon’s Terrestrial Animal Care Services (TeACS). The University of Oregon Institutional Animal Care and Use Committee (IACUC) reviewed and approved all care and experimental protocols.

### Songbird housing and acoustic recording

For experimental recordings, birds were singly housed in double-breeding cages with the divider removed to provide additional space (Model C-1510, Kings Cages International, Ft. Lauderdale, FL). Cages were placed inside Environmental Chambers (Omnitech Electronics Inc., Columbus, OH) lined with soundproofing foam to reduce external noise (Willtec Melamine SONEX Classic Noise Absorbing Foam Panels, Sonex, Minneapolis, MN). Fresh air was pumped into the chambers using tubes connected to a small air pump (Model #LPH80, JEMCO). Light cycle timing was adjusted to match the corresponding daylight hours in Eugene, Oregon. Enrichment for experimental birds included socialization (1-2 hours per day, 3-5 times per week) and daily treats and toys (dried mealworms, dried fruit, crinkle paper, and mixed greens).

We collected song recordings using an omnidirectional microphone (Audio-Technica AT803) positioned above each cage. We used an M-Audio 8 pre-amplifier to record each channel and route the recordings to the computer. Songs were detected and recorded using Sound Analysis Pro 11 software (Tchernichovski et al., 2000). Recordings were conducted at 44.1 kHz (single channel) and saved as WAV files. All recorded songs were undirected (sung in the absence of females and other male birds).

### Song detection and annotation

We isolated song segments from background noise using a supervised song detector trained on ground-truth labels marking song starts and stops, eliminating calls, cage noise, and extended silences from raw recordings (Vengrovski et al., 2026). We then annotated songs using a previously published, self-supervised song annotator that applies a masked-prediction vision transformer model to spectrograms (Vengrovski et al., 2026). TweetyBERT embeddings form distinct clusters corresponding to canary song syllables. Uniform Manifold Approximation and Projection (UMAP) was used for dimensionality reduction, followed by Hierarchical Density-Based Spatial Clustering with Noise (HDBSCAN) to identify syllable clusters (Vengrovski et al., 2026). Unlike the prior supervised model, TweetyNET (Cohen et al., 2022), this new supervised classifier is trained not on human labels but on labels automatically generated by TweetyBERT.

Because fitting UMAP is computationally and memory-intensive, the original TweetyBERT annotation pipeline was limited to approximately an hour of birdsong per bird. This sampling was enough to capture every syllable in a canary’s repertoire (Nottebohm et al., 1986; Vengrovski et al., 2026), and therefore sufficient for initial label discovery, but not enough to track longitudinal changes across thousands of songs. To scale beyond this limit, we developed a two-stage pseudolabeling pipeline. In the first stage, following the original TweetyBERT protocol, we obtained TweetyBERT embeddings from about 45 minutes of song, taking the same number of sampled time bins pre- and post-lesion. We reduced the pooled embeddings with UMAP and clustered them with HDBSCAN to assign each spectrogram time bin a bird-specific syllable label. In the second stage, we used these cluster assignments as pseudolabels to train a supervised decoder attached to TweetyBERT. The decoder maps the 196-dimensional attention output of the final transformer block through a three-layer Multilayer Perceptron (MLP) (196→128→64→C, with GELU activations), where C is the number of syllable states discovered for that bird. We jointly optimized this decoder and the TweetyBERT backbone using multiclass cross-entropy loss.

During inference, we normalized detected song segments, divided them into fixed-length contexts, and passed them through the network to produce a syllable-state prediction at each time bin. Changes between predicted states were then converted into syllable onset and offset annotations. This procedure reproduces the original HDBSCAN state assignments without applying UMAP to the full longitudinal dataset, allowing incremental processing of large datasets. Phrase duration was defined as the duration of consecutive time bins marked with a given syllable label (Fig.2B). To assess whether lesion-associated acoustic changes produced a gross reorganization of the embedding structure used to generate syllable labels, we visualized the pre-lesion and post-lesion portions of each bird’s UMAP embeddings (Supplemental Fig. 2). We used this comparison as a qualitative quality control.

### Lesion and sham lesion surgeries

Surgical protocols followed previously published lesion protocols targeting song nuclei in the Anterior Forebrain Pathway (AFP) (Goldberg and Fee, 2011; Koparkar et al., 2024). Birds were induced with 3-5% isoflurane (Isospire, Dechra Veterinary), anesthetized with Ketamine (Zetamine ketamine hydrochloride injection, VetOne) and Midazolam injection (USP), and maintained at 1-3% isoflurane for the remainder of the surgery. Birds were mounted in a stereotaxic device (Kopf), stabilized by non-rupture ear bars and a custom beak bar, and positioned so that the head angle of the anterior, flat portion of the skull was set to 20 degrees relative to the surface of the surgery table. Bilateral lesions were made by injecting N-methyl-D-aspartic acid (NMDA; Sigma, St. Louis, MO) to target Area X. We based our injection coordinates on previous lesions targeting Area X in zebra finches (Goldberg and Fee, 2011). We then adjusted injection volumes, numbers, and locations based on histology to better target Area X in canaries. The outer skull leaflet above the midsagittal sinus (“lambda”) was removed, and stereotaxic coordinates were set to zero at this point. Injections were typically between 5.40-5.60 mm anterior to lambda, 0.85-1.60 mm lateral to the skull’s midline, and 2.60-3.70 mm ventral to the surface of the brain. Sham lesions were made by injecting sterile saline solution (Hospira 0.9% sodium chloride injection, USP).

### Histology and lesion assessment

Birds were anesthetized with isoflurane and then euthanized with an overdose of pentobarbital (Euthasol, 150 mg/kg), shortly before a transcardial perfusion with 1× phosphate-buffered saline (Growcells.com MRGF-6396, diluted) for 4 minutes at a pump speed of 7.56 rpm, followed by fixation in 4% PFA diluted in 1× PBS for 10 minutes at 7.56 rpm (Rabbit-Plus Peristaltic Pump, Rainin). Brains were stored in 4% PFA overnight, then transferred to fresh 4% PFA for 24- 48 hours, then to 1× PBS for 1- 7 days before sectioning.

Brain tissues were embedded in 2.5% agarose (Sigma-Aldrich, AG6013-25G Type I, low EEO Agarose) mixed with 1× PBS. Brains were sectioned using a Leica VT 1200S vibratome into 50- or 75-μm-thick sections, mounted on slides (Fisherbrand Superfrost Plus Microscope Slides), and dried overnight. Slides were then rehydrated in a graded ethanol series, stained with diluted thionin solution, dehydrated in a graded ethanol series, cleared with Histoclear, and coverslipped with mounting media (Epredia Shandon-Mount). This protocol was based on other staining protocols for visualizing AFP song nuclei in seasonal songbirds (Smith et al., 1997; Singh et al., 2003), as well as a publicly available Nissl staining protocol (Huang, n.d.). We imaged slides at 5× magnification on a Leica Widefield Scope, and stitched the mosaic using LASX and Thunder software. The Nissl-defined borders of song nuclei and the striatal lamina were identified, and lateral or medial Area X lesion types were observed and noted qualitatively (Fig.1), with distinctions between medial and lateral based on the contours of the nucleus in the canary brain atlas (Vellema et al., 2011).

## Statistical Analysis

### Quantifying changes to phrase duration variability and syllable selection

We tested whether stuttering-like behavior differed among the three lesion groups: sham saline, birds with lateral-only AFP lesions, and birds with lesions to both the medial and lateral AFP (the “medial+lateral” group, pooling complete and partial medial+lateral lesion cases since both damaged medial AFP circuitry). As above, phrase duration is the length of a continuous stretch in which one syllable is repeated before the bird switches to a different syllable (Fig.2B, C). Supplemental Fig.1 shows example phrase duration distributions for early pre-lesion, late pre-lesion, and post-lesion in different birds within each lesion group.

We quantified phrase duration variability using the coefficient of variation (*CV*):

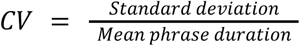

*CV* expresses variability in phrase duration, relative to the average phrase duration. Syllables were included if they contained at least 10 phrase occurrences from at least two recording days during both the late pre-lesion period (days -14 to -1) and the post-lesion period (days +1 to +14).

To prevent differences in phrase occurrence counts from biasing comparisons between the late pre-lesion and the post-lesion periods, we matched the number of observations separately for each syllable. The number of phrase occurrences sampled for each period was set to the smaller of the two available sample sizes (Supplemental Table 1). We randomly sampled this number of phrase occurrences without replacement from each period and calculated *CV* separately for the late-pre-lesion and post-lesion samples. We repeated this procedure 200 times, then averaged the resulting *CV* estimates across the 200 draws.

For each qualifying syllable, we calculated the change in phrase duration variability associated with the lesion as:

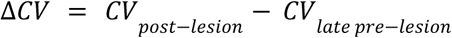

Positive Δ*CV* values indicate greater phrase duration variability after the lesion relative to the mean phrase duration.

Because the increase in phrase duration variability was concentrated in a subset of syllables (see example birds in Fig. 2C, Supplemental Figure 1), our primary analysis summarized the upper tail of each bird’s syllable-level Δ*CV* using its 75th percentile (Q75), yielding one independent Q75 Δ*CV* value per bird for between-group comparisons. Syllables with Δ*CV* values at or above the individual bird’s Q75 threshold (≥Q75) were plotted in Figure 3A and used for visualizing the time course of the effect and impacts on syllable acoustics.

**Figure 3.**
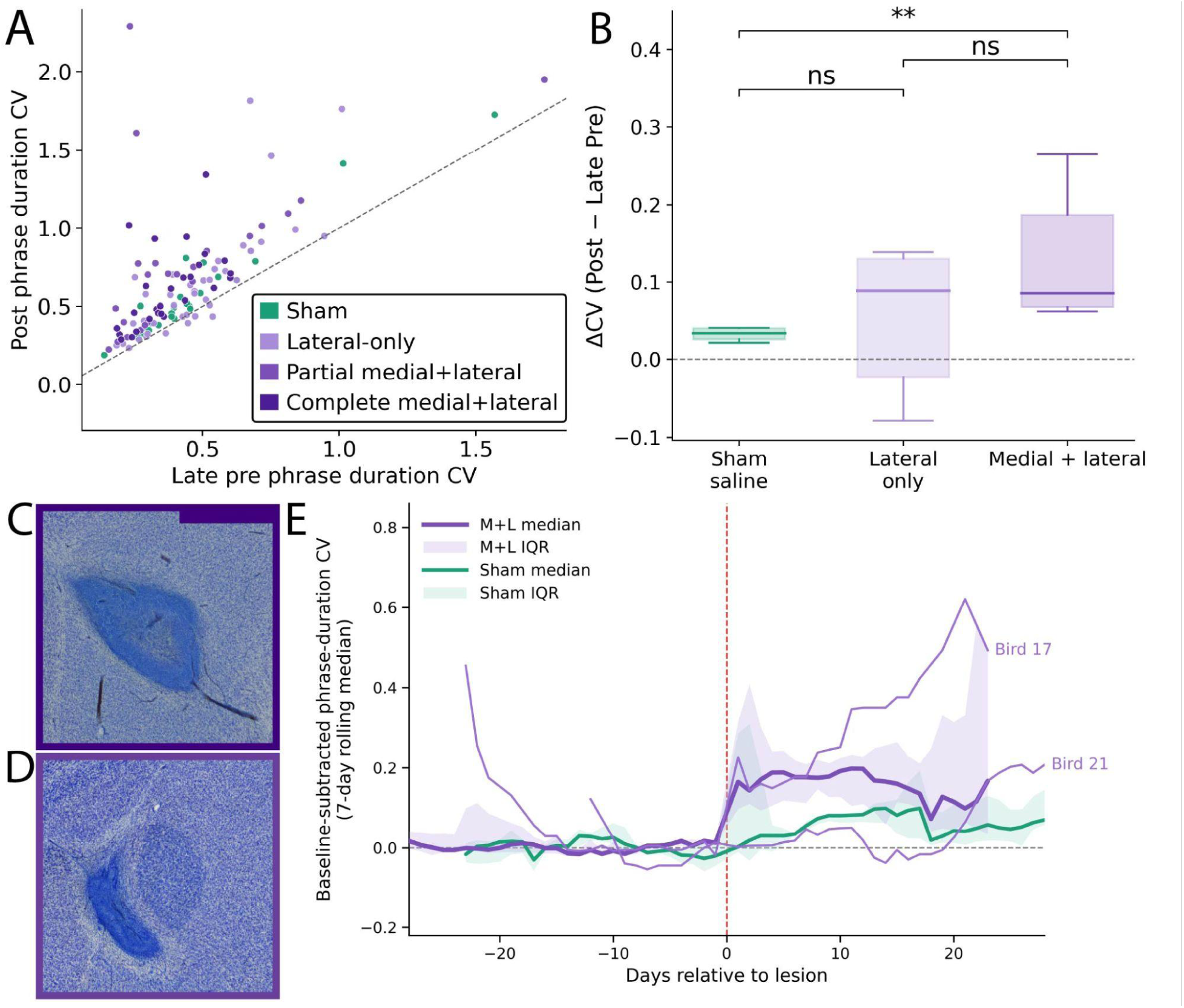
Lesions involving medial AFP circuitry produce a sustained increase in phrase duration variability. A) Late pre-lesion versus post-lesion phrase duration *CV* for syllables at or above each bird’s Q75 Δ*CV* threshold. The dashed line *y* = *x* indicates no change between periods. B) Q75 Δ*CV* averaged per bird and grouped by lesion group. Boxes show the median and interquartile range; whiskers span the 5th–95th percentiles. Group comparisons used exact permutation tests on differences in mean bird-level Q75 Δ*CV*, with Holm correction across the three planned comparisons. Medial+lateral lesion birds differed from sham controls (Holm-adjusted p = 0.0042), but not from lateral-only lesion birds (p = 0.1067); lateral-only lesion birds did not differ from sham controls (p = 0.6687). C–D) Representative Nissl-stained sections showing complete (C) and partial (D) medial+lateral AFP lesions. E) Time course of baseline-subtracted phrase duration *CV* for Q75 syllables. Thick lines and shading show the equal bird median and interquartile range for medial+lateral lesion birds (purple) and sham controls (teal). Lighter purple lines show representative medial+lateral lesion birds (Bird 17 and 21). We smoothed values using a centered 7-day rolling median. The vertical dashed line indicates the lesion date.

To test sensitivity to the Q75 threshold, we repeated the analysis using the 60th and 90th percentiles (Q60, Q90) of the same qualifying syllable-level distributions (Supplemental Table 2). No syllables were reselected across analyses. Q60, Q75, and Q90 are alternative summaries of the same qualifying syllables within each bird.

For each quantile, the permutation test statistic was the difference in mean bird-level quantile between lesion groups. We computed exact permutation p-values across all possible group assignments. Tests were one-sided (medial+lateral>sham; medial+lateral>lateral-only) or two-sided (lateral-only vs. sham), and Holm-corrected across three planned comparisons within each quantile.

### Comparing phrase duration variability between lesion groups

We compared bird-level Q75 Δ*CV* among lesion groups using exact permutation tests. The test statistic was the difference in mean bird-level Q75 Δ*CV* between groups. For each pairwise comparison, we enumerated all possible assignments of birds to the two groups, while preserving the observed group sizes. Because we predicted that the medial+lateral group would show the largest increase, we used a one-sided test to compare medial+lateral lesions against sham controls and lateral-only lesion birds, with the alternative hypothesis that Q75 Δ*CV* was greater in the medial+lateral group. We used a two-sided test to compare lateral-only lesion birds with sham controls. We Holm-corrected p-values across the three planned comparisons (Fig. 3B).

### Time course of post-lesion phrase duration variability

To examine whether the increase in phrase duration variability was concentrated immediately after the lesion surgery, or, instead, if it persisted across the post-lesion recording period, we examined the daily time course in birds with medial+lateral lesions and sham lesions (Fig.3E).

For each recording day, we calculated *CV* for each ≥Q75 syllable with at least 10 phrase occurrences. We then took the median across the syllables to obtain one daily *CV* value for each bird. To express daily variability relative to baseline, we subtracted each bird’s median late pre-lesion CV (days -14 to -1) from its daily *CV*:

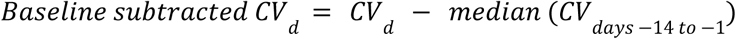

Where

*CV_d_* is the bird’s *CV* on recording day, *d*. A baseline-subtracted *CV* of 0 indicates no change from the bird’s late pre-lesion baseline. We smoothed each bird’s daily value using a centered 7-day rolling median. We calculated this separately for pre-lesion and post-lesion days so rolling windows did not span the lesion. For each day relative to the lesion, we then calculated the median and interquartile ranges across the birds. Each bird contributed one value per day, and therefore received equal weight. We plotted group summaries only on days with data from at least three birds.

To illustrate variability among individuals, we overlaid trajectories from two medial+lateral lesion birds (Bird 17 and 21; Supplemental Table 1) on the group summary in Fig.3E. Trajectories for all medial+lateral lesion and sham birds appear in Supplemental Figure 3.

### Quantifying AFP lesion-associated changes in syllable acoustic structure

We quantified the stability of syllable acoustic structure over time using the Bhattacharyya coefficient (*BC*), a general measure of overlap between two probability distributions (Bhattacharyya, 1943). Higher *BC* values indicate greater overlap between distributions and therefore greater stability of the represented acoustic structure.

For each syllable, we projected its TweetyBERT embeddings into UMAP space using fixed embedding parameters for each comparison. We then generated normalized density distributions from the projected embeddings, and calculated the *BC* between distributions from different recording periods. We split both pre-lesion and post-lesion recordings into early and late halves. The pre-lesion *BC* measured the overlap between the early and late pre-lesion halves, and the post-lesion *BC* measured overlap between the early and late post-lesion halves. This analysis compared the syllable acoustic structure stabilities pre- and post-lesion.

To control sample size for each syllable and avoid estimating acoustic distributions from a handful of longer renditions, we used a sampling procedure to balance run counts across the four comparison periods while keeping each run fully intact. Before identifying contiguous runs, we replaced noise labels with the nearest non-noise label. We applied temporal majority-vote smoothing using a 200-bin (∼540ms) window, which is used in syllable label smoothing for TweetyBERT. For each syllable cluster, we first identified contiguous phrase label “runs” separately within four periods: early pre-lesion, late pre-lesion, early post-lesion, and late post-lesion. A run qualified if it lasted at least 100ms and had at least 80% of its duration within its assigned period. A syllable cluster then qualified for analysis only if all four periods contained at least 15 qualifying runs. From each qualifying cluster, we randomly selected an equal number of runs per period, matching whichever period had the fewest qualifying runs. Syllables were retained for aggregate BC analysis only if the selected runs contained at least 10 seconds of acoustic data in every four periods.

We used all TweetyBERT time bins within those selected runs to calculate the Bhattacharyya coefficient (BC, see below). For each syllable, we pooled the embeddings from the selected runs across all four periods to define a shared two-dimensional UMAP density grid (20x20 bins, capturing 99% of points). We then calculated separate normalized density histograms for each period using this shared grid. For two normalized distributions, *p* and *q*, with probabilities *q* in UMAP bin *i*, *BC* was calculated as:

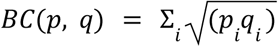

*BC* ranges from 0 to 1, with values closer to 1 indicating greater overlap.

For each bird, we took the median *BC* across its qualifying syllable clusters. Each bird contributed one paired pre/post value and carried equal weight regardless of the number of qualifying syllables. We again pooled complete and partial medial+lateral lesions because both involved damage to medial AFP circuitry. For each lesion group (sham, lateral–only, and medial+lateral), we compared pre- and post-lesion *BC* using two-sided paired Wilcoxon signed-rank tests. P-values were corrected across the three lesion-group comparisons using the Holm procedure (Fig. 4C). Summarizing to one value per bird avoids treating multiple syllables from the same bird as independent biological replicates.

**Figure 4.**
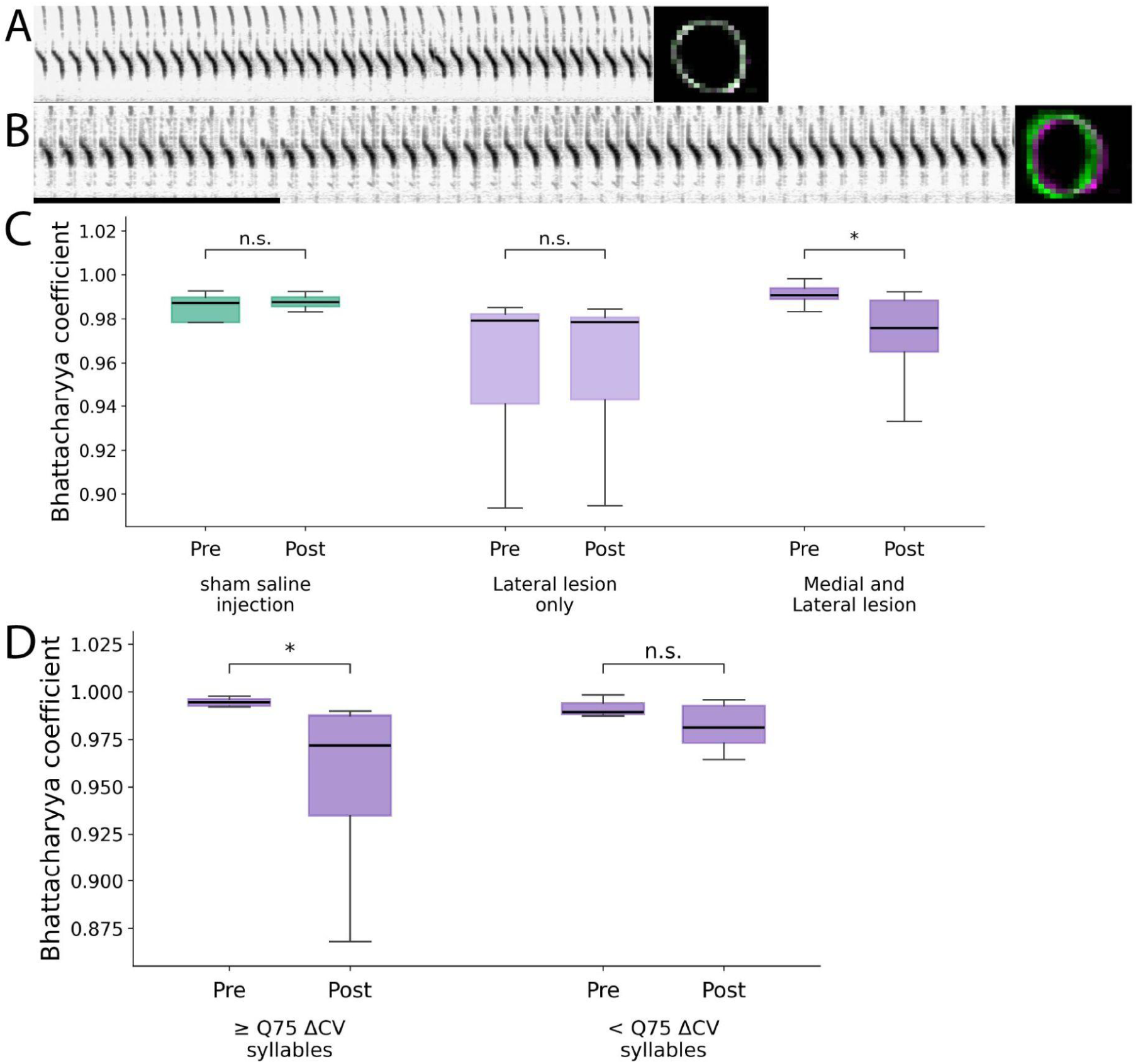
Lesions involving medial AFP circuitry are associated with a minor reduction in syllable acoustic stability. A) Example pre-lesion phrase spectrograms and overlap between early and late pre-lesion syllable embedding-space density distributions (*BC* = 0.995). B) Example post-lesion phrase spectrograms and density overlap for the same syllable cluster (*BC* = 0.623); scale bar, 1 s. This example was selected to illustrate a clearly visible change in embedding-space overlap, but it is substantially larger than the typical bird-level shift observed after medial+lateral lesions. C) Pre- and post-lesion Bhattacharyya coefficient (*BC*), summarized for each bird as the median across all qualifying syllable clusters. *BC* decreased after medial+lateral lesions (n = 8 birds; Holm-adjusted p = 0.0469), but not after lateral-only lesions (n = 8; p = 0.9453) or sham treatment (n = 4; p = 0.7500). D) Within medial+lateral lesion birds, syllables at or above each bird’s Figure 3 Q75 Δ*CV* threshold showed a significant post-lesion decrease in *BC* (n = 8; Holm-adjusted p = 0.0156), whereas syllables below Q75 did not (n = 8; p = 0.0781). P-values are from two-sided paired Wilcoxon signed-rank tests with Holm correction within each panel.

We next asked whether the drop in acoustic stability was concentrated among “stuttered” syllables. This analysis included only birds with medial+lateral lesions and used the same qualifying syllables and bird-specific Q75thresholds defined for the phrase duration analysis. Within each bird, qualifying *BC* clusters were divided into two groups based on the bird-specific Q75 Δ*CV* threshold. Clusters with Δ*CV* values at or above Q75 were classified as ≥Q75, whereas those below the threshold were classified as <Q75. For each group, we calculated the median pre-lesion and post-lesion *BC* across qualifying clusters within each bird. We then compared the paired pre- and post-lesion values using two-sided Wilcoxon signed-rank tests. *p* -values were corrected across the two comparisons using the Holm procedure (Fig. 4D). Birds were included in a comparison only if they contributed at least one qualifying *BC* cluster to that group.

Because *BC* compares early and late acoustic embedding distributions, a lower post-lesion *BC* could reflect greater short-term variability, gradual acoustic drift over time, or both. We therefore interpret lower *BC* as reduced stability rather than as a direct measure of acoustic variability alone.

## Results

### Histology

Although our injections targeted Area X, histology showed that some lesions extended into the surrounding nidopallium and damaged portions of LMAN and possibly MMAN. We therefore interpret our results as effects of lesions involving the broader AFP rather than Area X alone. Lesion location and extent varied across birds. Sham injections produced no detectable damage to Area X. Some lesions involved only lateral Area X, while others partially involved both medial and lateral Area X. In the most extensive cases, neither medial nor lateral Area X remained visible post-lesion. The position of these sections relative to the midline and surrounding landmarks supported classifying them as complete medial+lateral lesions. In adjacent sections, HVC, RA, and other structures remained visible, confirming that the absence of Area X reflected the lesion rather than poor staining or imaging.

Lesions were classified according to their histological relationship to medial and lateral Area X. Nissl-stained sections and surrounding anatomical landmarks were used to identify intact medial and lateral Area X (Fig.1B, C) and compared to the 3D canary brain atlas (Vellema et al., 2011), then used to classify birds as sham controls, lateral-only lesions, partial medial+lateral lesions, or complete medial+lateral lesions. Fig. 3C and D show representative complete and partial medial+lateral lesions. We used these categories in subsequent analyses of behavioral and acoustic effects.

### Lesions involving medial AFP circuitry increase phrase duration variability

AFP lesions produced a stuttering-like behavior in which birds varied more in the amount of time they spent repeating certain syllables before transitioning to the next syllable (Fig. 2B). This effect was not distributed evenly across the repertoire, but was concentrated in a subset of syllables (Fig. 2C; Supplemental Fig. 1).

To quantify these changes, we calculated Δ*CV* for each qualifying syllable as the post-lesion *CV* minus the late pre-lesion *CV*, so positive Δ*CV* values indicate greater phrase duration variability after lesion. For each bird, we used the 75th percentile (Q75) of the syllable Δ*CV* distribution to summarize the upper end of the observed changes. Syllables at or above each bird’s Q75 threshold are plotted in Fig. 3A, where the *y* = *x* identity line marks no change (points above the line increased in *CV* post-lesion, points below decreased). Because multiple syllables from the same bird are not independent biological replicates, we used one Q75 Δ*CV* value per bird for all between-group comparisons.

Birds with medial+lateral lesions had greater Q75 than sham controls (one-sided permutation test, Holm-adjusted p=0.0042; Fig. 3B). The medial+lateral group did not differ significantly from birds with lateral lesions only (p=0.1067), and birds with lateral lesions only did not differ significantly from sham controls (p=0.6687). Therefore, the clearest increase in phrase duration variability occurred in birds with lesions involving both medial and lateral AFP circuitry.

We tested whether this group pattern depended on using Q75 as the bird-level summary. The medial+lateral versus sham difference remained positive when each bird’s syllable-level Δ*CV* distributions were summarized at Q60, Q75, and Q90 (Supplemental Table 2). Statistical support was strongest at Q75 (one-sided exact label permutation test of difference in mean bird-level Q75 Δ*CV*, Holm-adjusted p=0.0042). The Q60 comparison did not reach significance (raw p = 0.0503, Holm-adjusted p = 0.1510), and the Q90 comparison was significant before but not after correction across the three lesion group comparisons (raw p = 0.0196, Holm-adjusted p = 0.0587). The direction of the medial+lateral versus sham effect was consistent across the upper-distribution summaries, but the strongest evidence was around Q75 rather than uniformly across all tested quantiles. These results indicate that the effect is located in the upper portion of the syllable-level response distribution, but that its detectability is sensitive to where the summary is taken in the upper range.

We next examined whether the increase in variability was limited to the period immediately after lesion or persisted over time. Because daily *CV* values were baseline-subtracted, 0 represents each bird’s median *CV* during the late prelesion period. In birds with medial+lateral lesions, the 7-day rolling median rose above baseline after lesion. It remained elevated across much of the post-lesion recording period. In contrast, the sham group median remained near baseline (Fig.3E). Individual medial+lateral lesion birds’ increases were variable, as illustrated by Birds 17 and 21 (Fig.3E; individual trajectories for all birds shown in Supplemental Figure 3). This descriptive time course suggests the effect was not confined to the immediate postoperative period.

Because phrase durations were calculated from TweetyBERT annotations, we also examined whether post-lesion song showed large changes in acoustic embedding structure used for syllable labeling. Across lesion groups, major bird-specific syllable clusters remained visually recognizable in paired pre-lesion and post-lesion UMAP plots (Supplemental Fig.2). We did not observe an obvious global collapse or reorganization of embedding structure post-lesion. This comparison provides qualitative control, rather than an independent measurement of decoder accuracy.

### Lesions involving medial AFP circuitry produced subtle but detectable changes to syllable acoustic structure

To determine whether lesions involving medial AFP circuitry also affected syllable acoustic stability, we measured changes in syllable distributions and their overlap in acoustic latent space. Greater overlap between the early and late embedding distributions of a syllable indicates greater acoustic stability across a recording period. We quantified this overlap using the Bhattacharyya coefficient (*BC*), comparing early versus late overlap before and after lesion (Fig.4A-C), and took the median *BC* across each bird’s qualifying syllables. Of the nine medial+lateral lesion birds included in the phrase duration *CV* analysis, eight contributed to the *BC* analysis (Bird 20 was excluded because no syllable cluster met the four-period data requirement; see Supplemental Table 1).

Birds with complete or partial medial+lateral AFP lesions showed a small but significant reduction in *BC* after lesion, that is, reduced overlap between early and late syllable embedding distributions (n = 8 birds; median prelesion *BC* = 0.9907, median postlesion *BC* = 0.9758, median paired Δ*BC* = −0.0115; two-sided paired Wilcoxon signed-rank test, Holm-adjusted p=0.0469; Fig. 4C). Here, Δ*BC* was calculated as postlesion *BC* minus prelesion *BC*, so negative values indicate reduced acoustic stability after lesion. These results indicate that medial+lateral AFP lesions were associated with lower embedding space overlap across the postlesion recording period.

Sham controls showed no significant change in *BC* (n = 4 birds; median prelesion *BC* = 0.9871, median postlesion *BC* = 0.9876, median paired Δ*BC* = +0.0022; Holm-adjusted p=0.7500; Fig. 4C). Birds with lateral lesions only also showed no significant change (n = 8 birds; median prelesion *BC* = 0.9791, median postlesion *BC* = 0.9785, median paired Δ*BC* = −0.0003; Holm-adjusted p=0.9453). Thus, the clearest reduction in acoustic stability across all qualifying syllable clusters occurred in birds with medial+lateral AFP involvement.

We next asked whether the change in acoustic stability was concentrated among stuttered syllables. Within birds with medial+lateral lesions, we classified qualifying syllables using the same bird-specific Q75thresholds used in the phrase duration analysis. Stuttered syllables, defined as those at or above Q75, showed a significant reduction in *BC* after lesion (n = 8 birds; median prelesion *BC* = 0.9945, median postlesion *BC* = 0.9718, median paired Δ*BC* = −0.0192; two-sided paired Wilcoxon signed-rank test, Holm-adjusted p=0.0156; Fig. 4D). In contrast, syllables below Q75 did not show a significant change (n = 8 birds; median prelesion *BC* = 0.9892, median postlesion *BC* = 0.9813, median paired Δ*BC* = −0.0042; Holm-adjusted p=0.0781). Although the reduction among Q75 syllables was statistically detectable, its magnitude was small.

Overall, lesions involving medial AFP circuitry were associated with reduced embedding space overlap, and this reduction was most apparent among stuttered syllables. Because reduced *BC* may reflect increased occurrence-to-occurrence variation, progressive acoustic drift over time, or both, these results indicate a subtle lesion-associated change in syllable acoustic structure but do not distinguish between these possibilities.

## Discussion

### Stuttering-like increase in phrase duration variability

When lesions involved both the medial and lateral portions of the Anterior Forebrain Pathway (AFP), adult canaries developed a stuttering-like behavior: the time they spent repeating a syllable became more variable before they transitioned to the next syllable. This effect was concentrated in a subset of syllables rather than distributed uniformly across each bird’s repertoire. Birds with medial and lateral lesions had a greater increase in phrase duration coefficient of variation (*CV*) than sham controls. In contrast, birds with lateral-only lesions did not differ significantly from sham controls (Fig.3B). These findings implicate medial AFP circuitry in controlling phrase transitions during adult breeding-season canary song.

This increased variability lasted through much of the post-lesion period, rather than fading shortly after surgery. This time course differs from other lesioning studies in different songbird species. Partial Area X lesions in Bengalese finches produced temporary stuttering-like behavior that peaked shortly after surgery and resolved within three weeks (Kobayashi et al., 2001). In zebra finches, excitotoxic lesions of lateral Area X produced increased syllable repetition in a subset of birds, and the behavior appeared to worsen as Area X recovered after the lesion (Kubikova et al., 2014). By comparison, phrase duration variability in our canaries remained elevated throughout much of the post-lesion period. It is unclear if these differences are due to species-specific song organization or pre-lesion structure of repeated syllables.

The clearest behavioral effect occurred in birds with lesions involving both medial and lateral AFP. This pattern broadly matches prior work in Bengalese finches, which found no clear relationship between lesion size and stuttering-like behavior (Kobayashi et al., 2001). In our study, medial and lateral lesion birds showed greater increases in phrase duration than sham controls, whereas lateral-only lesioned birds did not differ significantly from sham controls. However, the medial and the lateral-only groups did not differ significantly from one another, and the medial damage never occurred without lateral damage in our birds. These experiments therefore implicate lesions involving medial AFP circuitry, but cannot isolate the independent contributions of medial AFP from lateral AFP.

These results may help interpret earlier Area X lesion studies. Kobayashi and colleagues did not distinguish medial from lateral Area X involvement (Kobayashi et al., 2001), so varying medial AFP damage across their subjects may explain the lack of clear relationship between lesion volume and behavioral impact. This is possibly consistent with other work finding that another nucleus in medial AFP, MMAN, contributes to adult Bengalese finch song (Koparkar et al., 2024). Similarly, an earlier Area X lesion study in adult canaries reported no observable song effects (Nottebohm et al., 1976). Interestingly, the published histology appears to show lesions concentrated in lateral Area X (Nottebohm et al., 1976). Together, these observations suggest that lesion location within the AFP may predict its effect size on adult birdsong.

We use “stuttering-like” descriptively to refer to prolonged and variable repetition of a syllable before transition, rather than to imply direct equivalence with developmental human stuttering. With that caveat, the behavior shares features with timing abnormalities observed in people who stutter. Adults who stutter exhibit increased variability in rhythmic timing and altered speech and nonspeech movement kinematics (Max and Yudman, 2003; Max et al., 2003; Daliri et al., 2014). Stuttering has also been associated with unstable sensorimotor integration and increased variability in speech movements (Olander et al., 2010; Sares et al., 2018; Wiltshire et al., 2021). Dopaminergic signaling within basal ganglia circuits has been implicated in human stuttering (Wu et al., 1997; Chang and Guenther, 2019), consistent with the involvement of Area X, a songbird basal ganglia nucleus containing medium spiny neurons (Ding and Perkel, 2002; Sasaki et al., 2006). There still remain important differences between the behavior we observed. Human stuttering typically begins in childhood and often remits, whereas the present effect was induced in adult canaries with fully learned, crystallized song and persisted across much of the post-lesion recording period (Reilly et al., 2013; Franken et al., 2018). Human stuttering also tends to occur at the start of words or utterances (Natke et al., 2004). In contrast, the increased variability here occurred across syllables in the repertoire and was not concentrated on the first syllable of the song. Despite these differences, the shared reliance on basal ganglia-dependent timing and sequencing makes the medial AFP a useful system for studying how circuits stabilize repeat motor elements.

### Detectable changes in acoustic stability

These results align with the broader view that the AFP may continue to play a role in producing crystallized adult song, specifically by contributing to variability, plasticity, and sequence regulation. Prior work found that lesions to Area X in adult male zebra finches reduced within-syllable variation (Kojima et al., 2018), and LMAN lesions can stabilize adult song after deafening, suggesting that AFP output contributes to ongoing song variability and experience-dependent modification, rather than song execution (Brainard and Doupe, 2001). A previous study in adult canaries found that LMAN lesions performed in fall reduced acoustic variance in fall plastic song. In contrast, lesions to LMAN during the breeding season had no apparent changes to syllable acoustics (Alliende et al., 2017). Similarly, LMAN lesions in adult Bengalese finches reduce syllable-structure variability (Hampton et al., 2009), supporting a role for AFP output in regulating acoustic variability. In our study, conducted during the canary breeding season, we found little change in sham and lateral-only lesion birds.

In contrast, medial and lateral AFP lesions showed a small but significant reduction in post-lesion acoustic embedding space overlap. The small scale of this effect could reflect changes in physiological demands during long stuttered bouts, such as fatigue in the respiratory muscles or the laryngeal musculature. This interpretation is consistent with the observation that the reduction reached significance only among stuttered syllables, although we did not test the difference between stuttered and non-stuttered syllables directly. Further investigation is needed to determine whether this minor change in variability reflects changes in peripheral motor planning or the AFP’s role in driving acoustic variability during song learning.

### Limitations

Although we performed histological classification blind to the behavioral results, we defined lesion groups post hoc from histology, so group membership is partly confounded with lesion extent. Medial damage never occurred without lateral damage. Group sizes were small (n = 4 sham, 8 lateral-only, 9 medial+lateral), and several comparisons had limited resolution: the four-bird sham group cannot reach significance in a within-group Wilcoxon test. Non-significant results here indicate absence of evidence rather than evidence of absence, including the fact that the two lesion groups did not differ significantly from one another. Several analytic choices (the Q75 summary, the ±14-day windows, the inclusion criteria, and the pooling of complete and partial lesions) were not pre-specified; we therefore regard these results as exploratory. Phrase boundaries were derived from a supervised decoder trained on pseudolabels spanning both pre- and post-lesion recordings; post-lesion song fell within its training distribution, and the lesion-associated acoustic change was very small in absolute terms; however, we did not formally quantify annotation accuracy by period.

### Conclusion and future directions

Our findings demonstrate that the medial AFP influences the phrase structure of adult breeding-season canary song. We performed bilateral excitotoxic lesions targeting the medial and lateral AFP circuits during the breeding season. We developed a novel analysis method to efficiently process thousands of canary songs. We found that lesions involving the medial AFP produced a sustained stuttering-like behavior in a subset of syllables, characterized by increased variability in syllable repeat time. Lesions to only the lateral AFP did not produce a detectable effect relative to sham. However, the two lesion groups did not differ significantly from each other, suggesting that the medial basal ganglia-thalamocortical circuit may contribute to the ongoing maintenance of phrase-level timing in crystallized song.

The medial AFP offers a compact, well-defined circuit for studying the neural basis of motor sequencing disorders, such as stuttering, in which causal intervention, higher-resolution behavioral quantification, and single-unit physiology can be combined.

## Code Accessibility

Analysis code is available at https://github.com/hulseyvincentr/AFP_lesion_paper. The combined TweetyBERT and high-throughput MLP song annotation pipeline is available at https://github.com/georgevenven/tweety_bert. The TweetyNET song detector is available at https://github.com/georgevenven/tweety_net_song_detector.

## Data Availability

Analysis-ready data supporting this study, including TweetyBERT song annotation files and processed data used in analyses, have been prepared for deposition in Zenodo and will be made publicly available upon publication at https://doi.org/10.5281/zenodo.22715762. During peer review, these data are available from the authors upon reasonable request. Additional raw data are available from the authors upon reasonable request because the complete datasets are too large to deposit in the repository.

## Author contributions

M.R.H.-V. and T.J.G. designed the research; M.R.H.-V. and E.S. performed the research; M.R.H.-V., G.V., and E.S. contributed unpublished reagents/analytic tools; M.R.H.-V. analyzed data; M.R.H.-V., G.V., E.S., and T.J.G. wrote the paper.

## Declaration of generative AI and AI-assisted technologies in the writing process

The authors used ChatGPT and Claude to assist with code development and to rephrase, simplify, and improve the manuscript’s prose and grammar. The authors reviewed, tested, and approved all code and text suggestions before submission.

