## Supplemental Materials for "Lesions Involving Medial Anterior Forebrain Pathway Circuitry Destabilize Phrase Timing in Adult Canary Song"

**Supplemental Table 1. Animal metadata and sample sizes for phrase duration analyses.** Each row represents one bird. Qualifying syllables had at least 10 phrase occurrences from at least two recording days in both the late pre-lesion period (days −14 to −1) and the post-lesion period (days +1 to +14). For each qualifying syllable, we matched the number of phrase occurrences in late pre-lesion and post-lesion periods by subsampling each period to the smaller sample size available, then calculated the balanced coefficient of variance ($CV$). We repeated this balancing 200 times and averaged estimates across draws. We calculated $\Delta CV$ as the post-lesion $CV$ minus the late pre-lesion $CV$. ≥Q75 syllables are syllables that were at or above the bird's Q75 threshold for $CV$. Balanced phrase observation per period is the number of phrase occurrences that contributed per period for the ≥Q75 syllables.

| Bird | Treatment date | Lesion group | Recorded days pre | Recorded days post | Qualifying syllables | ≥Q75 syllables | Balanced phrase observations per period (≥Q75) |
|---|---|---|---|---|---|---|---|
| 1 | 2024-03-07 | Sham | 16 | 34 | 24 | 6 | 2140 |
| 2 | 2024-03-05 | Sham | 25 | 34 | 28 | 7 | 5626 |
| 3 | 2025-03-17 | Sham | 57 | 18 | 21 | 6 | 3329 |
| 4 | 2025-03-04 | Sham | 19 | 24 | 18 | 5 | 5540 |
| 5 | 2025-05-22 | Lateral-only | 33 | 17 | 20 | 5 | 1374 |
| 6 | 2025-05-23 | Lateral-only | 29 | 7 | 14 | 4 | 231 |
| 7 | 2025-05-16 | Lateral-only | 22 | 12 | 12 | 3 | 1542 |
| 8 | 2024-01-23 | Lateral-only | 44 | 13 | 27 | 7 | 4298 |
| 9 | 2024-01-20 | Lateral-only | 35 | 26 | 38 | 10 | 5169 |
| 10 | 2025-03-18 | Lateral-only | 39 | 30 | 11 | 3 | 2352 |
| 11 | 2025-02-20 | Lateral-only | 34 | 22 | 23 | 6 | 5638 |
| 12 | 2025-03-20 | Lateral-only | 34 | 24 | 13 | 4 | 1079 |
| 13 | 2024-04-09 | Partial medial+lateral | 36 | 14 | 26 | 7 | 7145 |
| 14 | 2024-02-13 | Partial medial+lateral | 62 | 18 | 31 | 8 | 4846 |
| 15 | 2024-02-20 | Partial medial+lateral | 32 | 18 | 30 | 8 | 2833 |
| 16 | 2024-04-10 | Complete medial+lateral | 37 | 12 | 28 | 7 | 6564 |
| 17 | 2024-04-16 | Complete medial+lateral | 34 | 22 | 28 | 7 | 4281 |
| 18 | 2024-04-20 | Complete medial+lateral | 9 | 29 | 20 | 5 | 7403 |
| 19 | 2024-06-28 | Complete medial+lateral | 30 | 15 | 11 | 3 | 2255 |
| 20* | 2024-07-02 | Complete medial+lateral | 39 | 3 | 6 | 2 | 39 |
| 21 | 2024-05-01 | Complete medial+lateral | 20 | 17 | 12 | 3 | 2865 |

*Bird 20's limited post-lesion sample size coincided with the seasonal transition into molt, when canaries stop singing. He was included in the phrase duration analyses but excluded from the Bhattacharyya Coefficient analyses because no syllable cluster contained sufficient data in all four BC comparison periods.

**Supplemental Table 2. Sensitivity of lesion-group differences in phrase-duration variability to the bird-level quantile summary.**

To assess whether the primary Q75 result depended on the selected percentile, we repeated the analysis using the 60th, 75th, and 90th percentiles of each bird's distribution of syllable-level ΔCV (post-lesion CV − late pre-lesion CV). Values shown for each lesion group are bird-level summary statistics. Group comparisons used exact permutation tests with bird as the independent experimental unit. The test statistic was the difference in mean bird-level quantile ΔCV between groups. Tests comparing medial+lateral lesions with sham and lateral-only lesions were one-sided in the predicted direction (medial+lateral > comparison group); lateral-only versus sham was two-sided. We Holm-corrected p-values across the three planned lesion-group comparisons separately within each quantile. The medial+lateral versus sham effect was positive at Q60, Q75, and Q90, with the strongest statistical support at Q75.

| Quantile | Sham median ΔCV | Lateral-only median ΔCV | Medial+lateral median ΔCV | M+L vs Sham Holm-adjusted p | M+L vs Lateral-only Holm-adjusted p | Lateral-only vs Sham Holm-adjusted p |
|---|---|---|---|---|---|---|
| Q60 | -0.008 | 0.035 | 0.045 | 0.1510 | 0.1853 | 0.7374 |
| Q75 | 0.033 | 0.088 | 0.085 | 0.0042 | 0.1067 | 0.6687 |
| Q90 | 0.091 | 0.146 | 0.218 | 0.0587 | 0.0587 | 0.7071 |

Note. Bird is the independent biological replicate. The permutation-test statistic was the difference in mean bird-level quantile ΔCV between groups. M+L vs Sham and M+L vs Lateral-only were one-sided tests with the alternative M+L > comparison group; Lateral-only vs Sham was two-sided. We used exact bird-label permutations whenever feasible. Sample sizes: Sham n=4, Lateral-only n=8, Medial+lateral n=9.

**Detailed exact permutation results**

| Quantile | Comparison | Alternative | Mean difference | Raw p | Holm p within quantile | Holm p across Q60/Q75/Q90 |
|---|---|---|---|---|---|---|
| Q60 | Lateral-only vs Sham | two-sided | 0.014 | 0.7374 | 0.7374 | 1.0000 |
| Q60 | M+L vs Lateral-only | > | 0.056 | 0.0926 | 0.1853 | 0.1067 |
| Q60 | M+L vs Sham | > | 0.071 | 0.0503 | 0.1510 | 0.0503 |
| Q75 | Lateral-only vs Sham | two-sided | 0.021 | 0.6687 | 0.6687 | 1.0000 |
| Q75 | M+L vs Lateral-only | > | 0.075 | 0.0534 | 0.1067 | 0.1067 |
| Q75 | M+L vs Sham | > | 0.096 | 0.0014 | 0.0042 | 0.0042 |
| Q90 | Lateral-only vs Sham | two-sided | 0.020 | 0.7071 | 0.7071 | 1.0000 |
| Q90 | M+L vs Lateral-only | > | 0.131 | 0.0237 | 0.0587 | 0.0712 |
| Q90 | M+L vs Sham | > | 0.151 | 0.0196 | 0.0587 | 0.0392 |

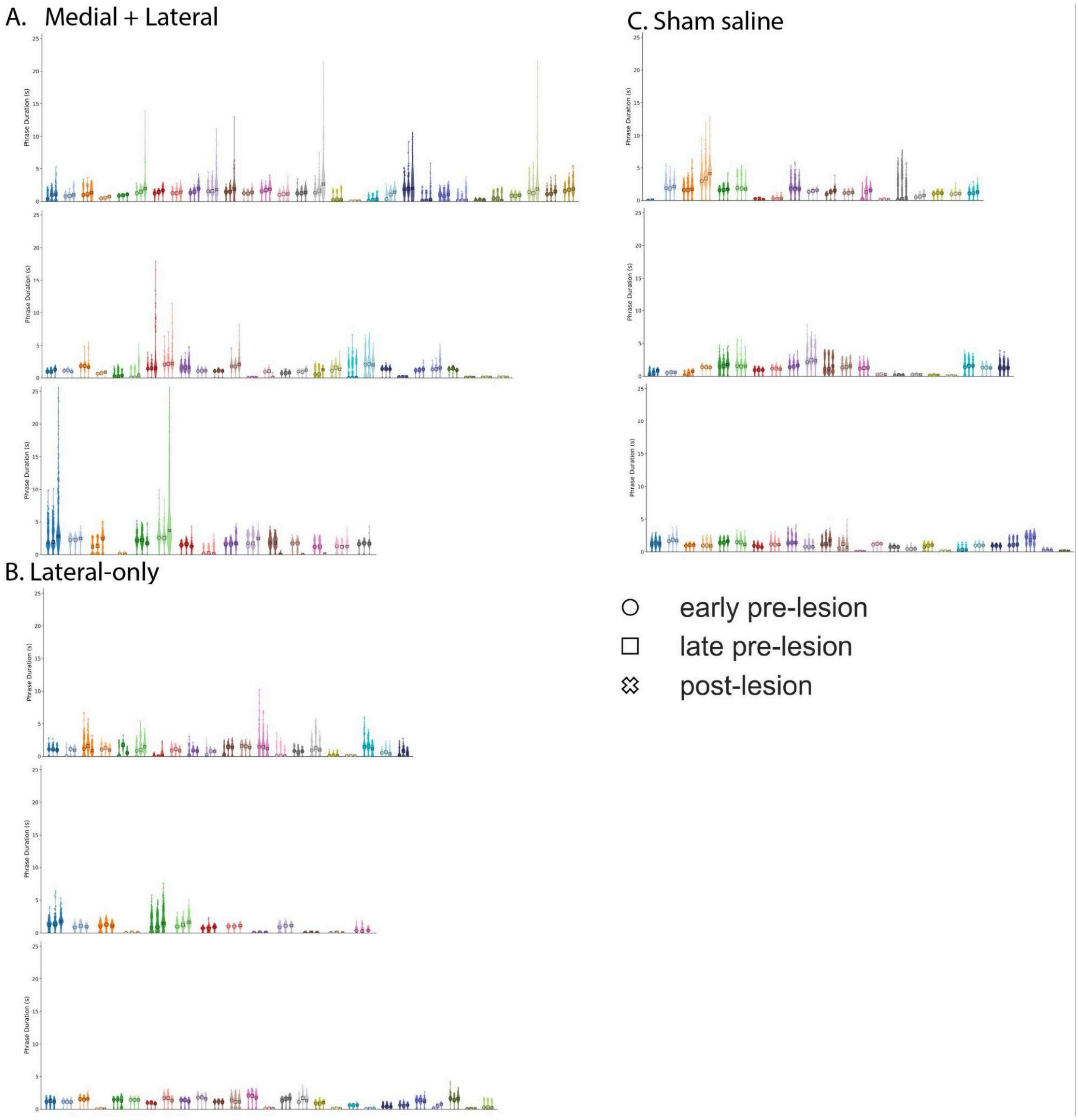


**Supplemental Figure 1. Phrase duration distributions by individual bird and lesion group.** Individual birds with A) medial+lateral AFP lesions (top to bottom: Birds 15, 16, and 21), B) lateral-only AFP lesions (Birds 5, 12, and 8), and C) sham saline injections (Birds 4, 3, and 1). Each row represents one bird; colors denote bird-specific syllable classes. Distributions are graphed for early pre-lesion, late pre-lesion, and post-lesion recording periods. All panels share a common 0- 25-second y-axis. Post-lesion changes were variable across birds and typically affected only a subset of syllable classes rather than the full repertoire.

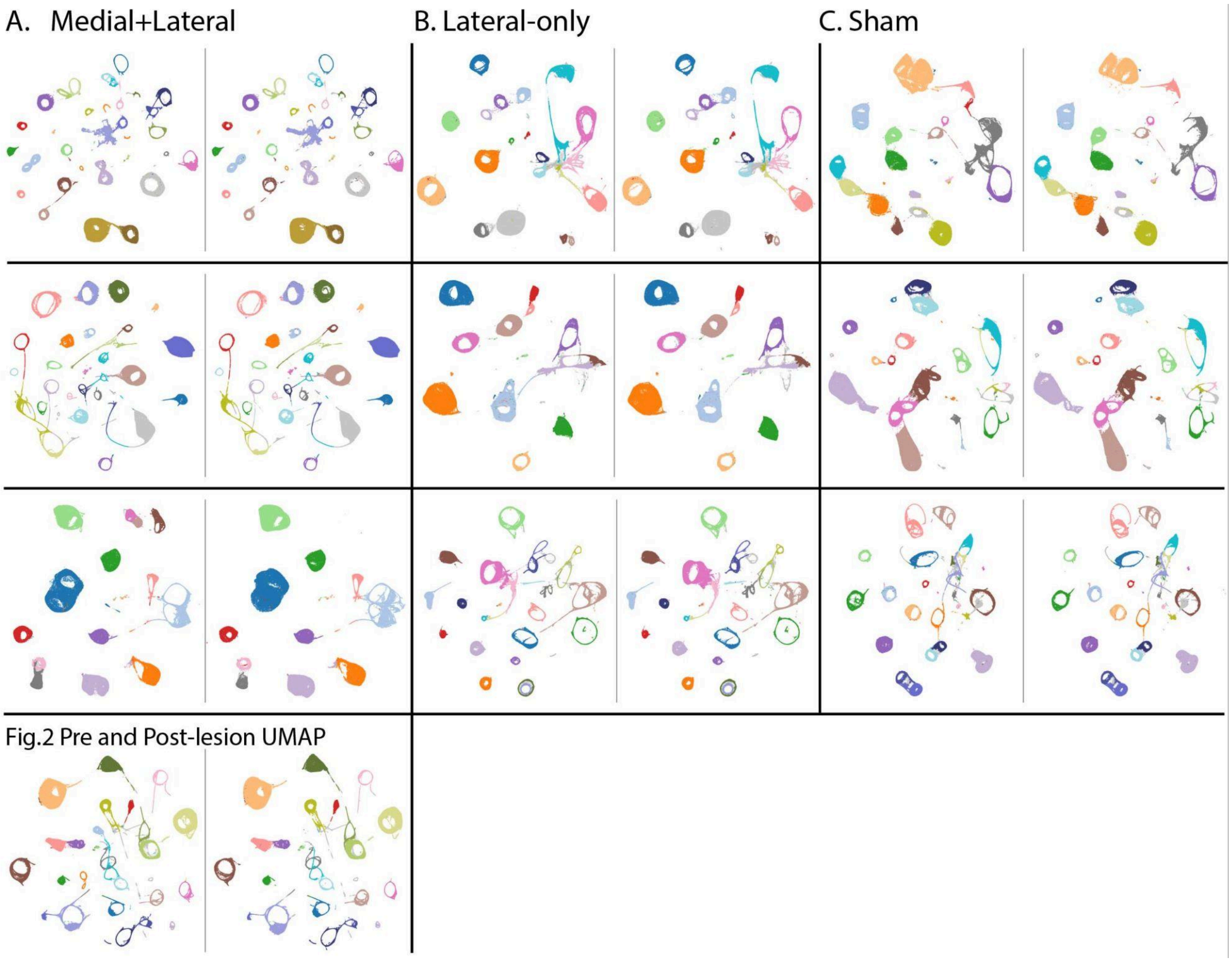


**Supplemental Figure 2. TweetyBERT embedding structure is qualitatively preserved across lesion groups.** Paired pre-lesion (left) and post-lesion (right) UMAP visualizations for representative birds from each lesion group. A) Medial+lateeral AFP lesions. The top three rows show the same birds as Supplemntal Fig.1 (top to bottom: Birds 15, 16 and 21); the bottom row shows Bird 13, the bird featured in Figure 2. B) Lateral-only AFP lesions (Birds 5, 12, 8). C) Sham saline injections (Birds 4, 3, 1). Each row shows one bird, with pre- and post-lesion points plotted in the same bird-specific UMAP space. Colors indicate the same bird-specific TweetyBERT/HDBSCAN syllable classes across periods. Major syllable clusters remained identifiable after lesion in all three groups, providing a qualitative check against large-scale reorganization of the embedding space that would confound syllable labeling.

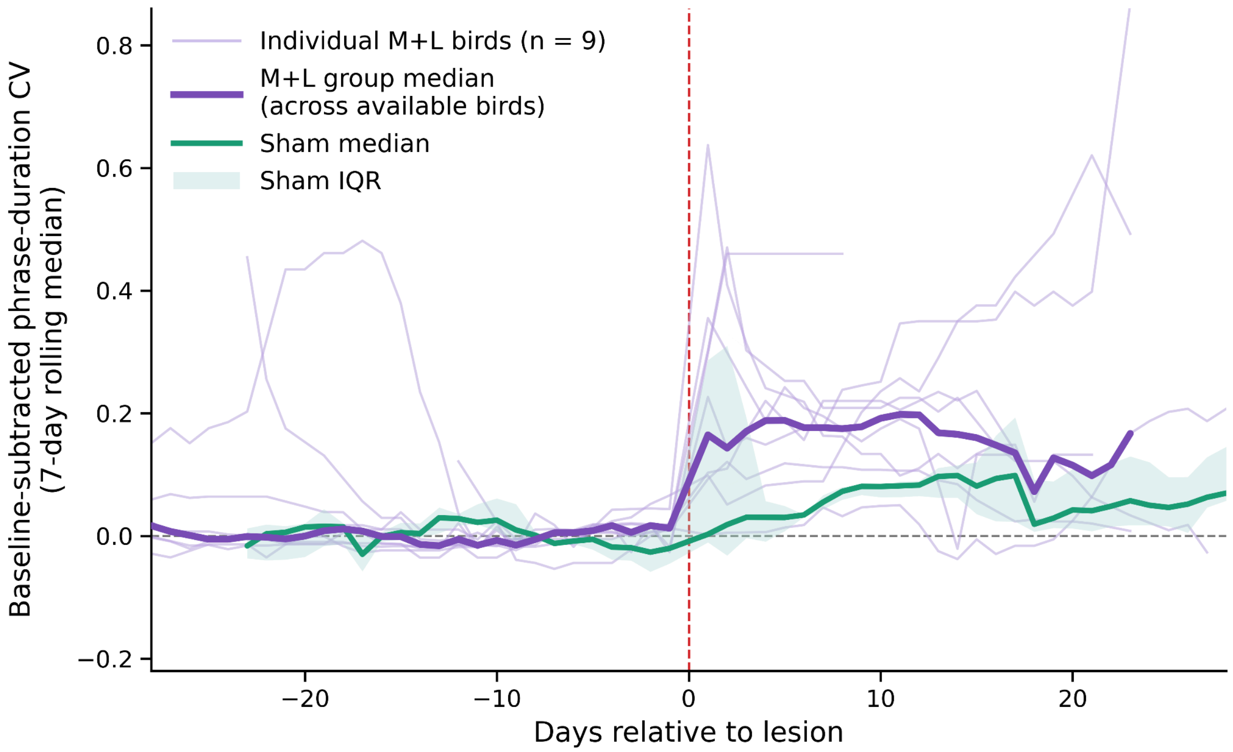


**Supplemental Figure 3. Individual time courses of phrase duration variability of medial+lateral AFP lesions.** Light purple lines show baseline-subtracted phrase-duration CV across days relative to lesion for each medial+lateral lesion bird (n=9); the thick purple lines show the median across these birds. The teal line shows the sham group median, with shading indicating the interquartile range across sham birds. For both groups, daily values reflect syllables at or above each bird's Q75 $\Delta CV$ threshold, smoothed with a centered 7-day rolling median. The vertical red dashed line marks the lesion date.
